# Hunger Recruits a Parallel Circuit Encoding Alcohol Reward

**DOI:** 10.1101/2025.10.14.682140

**Authors:** Kavin M. Nunez, Lewis M. Sherer, Anthony Walley, Sarah Salamon, Vivian M. Chan, Mustafa Talay, Gilad Barnea, Karla R. Kaun

## Abstract

Internal states like hunger, pain, thirst and arousal can bias behavior by affecting sensory and memory processing. Internal states are critical to understand in the context of alcohol addiction because they influence cravings, reinstatement, and relapse. Norepinephrine plays a key role in both hunger and alcohol-induced arousal and preference, but the circuit-level mechanisms through which it modulates the influence of hunger on alcohol preference are not well understood. We sought to address this using intersectional genetic tools for manipulating neurons expressing octopamine, the invertebrate analogue of vertebrate norepinephrine. We identified a single octopamine neuron required for ethanol seeking only when *Drosophila* are food-deprived. Hunger increased baseline activity in this neuron, making it more responsive to an odor cue previously paired with ethanol. A combination of genetic and connectome analyses revealed that synaptic partners of this octopaminergic neuron form a functional module that acts on *Drosophila* memory circuitry. Thus, we show that hunger recruits a parallel circuit that drives learned ethanol preference, providing a neuronal framework through which internal state influences the expression of memory for ethanol-associated cues.

## Introduction

The perception of internal sensations and states within the body, defined as interoception, influences the decisions animals make^1–5^. Psychoactive substances such as alcohol and drugs of abuse induce a unique interoceptive experience that interacts in complex ways with other internal states^6–9^. For example, hunger increases ethanol consumption through highly conserved, overlapping peptidergic, orexigenic and monoaminergic pathways^10–18^ in rats^19–22^, mice^23^, guinea pigs^24^, hamsters^25^, non-human primates^26^, and humans^27–29^. However, the mechanisms through which hunger influences the way ethanol reward is encoded and remembered are not well understood.

Norepinephrine is one of the best-characterized neurotransmitters underlying the effects of hunger on the brain^30–36^ and is associated with increases in alcohol-induced arousal and preference^37–41^. This makes norepinephrine an ideal candidate to investigate how hunger influences ethanol preference. Addressing this question within the mammalian noradrenergic system has been difficult due to the complex, heterogeneous, and broadly innervating connectivity that characterizes noradrenergic neurons in the central nervous system^42–51^. Furthermore, while norepinephrine plays important roles in both appetite regulation^52–58^ and alcohol consumption^59–68^, whether it directly influences how hunger alters alcohol consumption is unclear.

The invertebrate functional analogue of norepinephrine, octopamine, has been well-characterized^69–73^, making *Drosophila* an ideal model system to investigate this question due to the availability of intersectional genetic tools that provide high spatial resolution for manipulating individual octopaminergic cells within the brain^69,74–76^. Moreover, hunger can be tightly controlled and plays a key role in the plasticity of reward and memory circuitry^71,72,77–90^. While hunger influences the activity of octopamine neurons, how this influence alters the rewarding properties of ethanol is unknown. We hypothesized that hunger influences ethanol preference through octopaminergic circuitry. Through a combination of genetics, behavior, transsynaptic labeling, electron microscopy (EM) neuron reconstruction, and *in vivo* calcium imaging, we reveal a parallel octopaminergic circuit motif through which hunger facilitates cue-induced ethanol seeking.

## Results and Discussion

### Octopamine mediates hunger state-dependent ethanol memory expression

We trained flies to associate an olfactory cue with a rewarding dose of ethanol^91,92^ while satiated or hungry (Figure 1A and Figure S1A). Wild type control flies showed a characteristic shift from initial aversion to the ethanol associated cue, followed by approach over the course of 24 hours whether they were satiated or food-deprived (Figure S1).To investigate octopamine’s role in the ability to form a state-dependent memory for ethanol, we tested how disrupting octopamine synthesis or release impacted preference for the ethanol-associated cue 24 hours after training. Wild-type control flies show memory both when satiated and food-deprived, while tyramine-beta hydroxylase (Tβh)^nM18^ mutant flies that do not synthesize octopamine^93^ show reduced ethanol memory only when food-deprived (Figure 1B, 1C). We next tested whether synaptic transmission from octopamine/tyramine-producing neurons was required for odor-ethanol memory in satiated or hungry flies. Acutely blocking synaptic transmission in *Tyrosine decarboxylase 2* (*Tdc2*) neurons using a temperature-sensitive dominant negative form of dynamin, *shibire^ts^*^1^ ^94^, resulted in reduced ethanol memory in food-deprived, but not satiated flies (Figure 1D and Figure S2). Together, these observations suggest that one or more octopamine/ tyramine-producing neurons encode a “hunger” internal state that elicits cue-induced ethanol seeking in an octopamine-dependent manner. This is consistent with previous research showing a state-dependent response of octopamine/ tyramine signaling in foraging or feeding^81,83,85,88,95–98^, sensory processing^78,85^, memory^69,70,99–101^, metabolic traits, and starvation resistance^71,72,102,103^.

**Figure 1.**
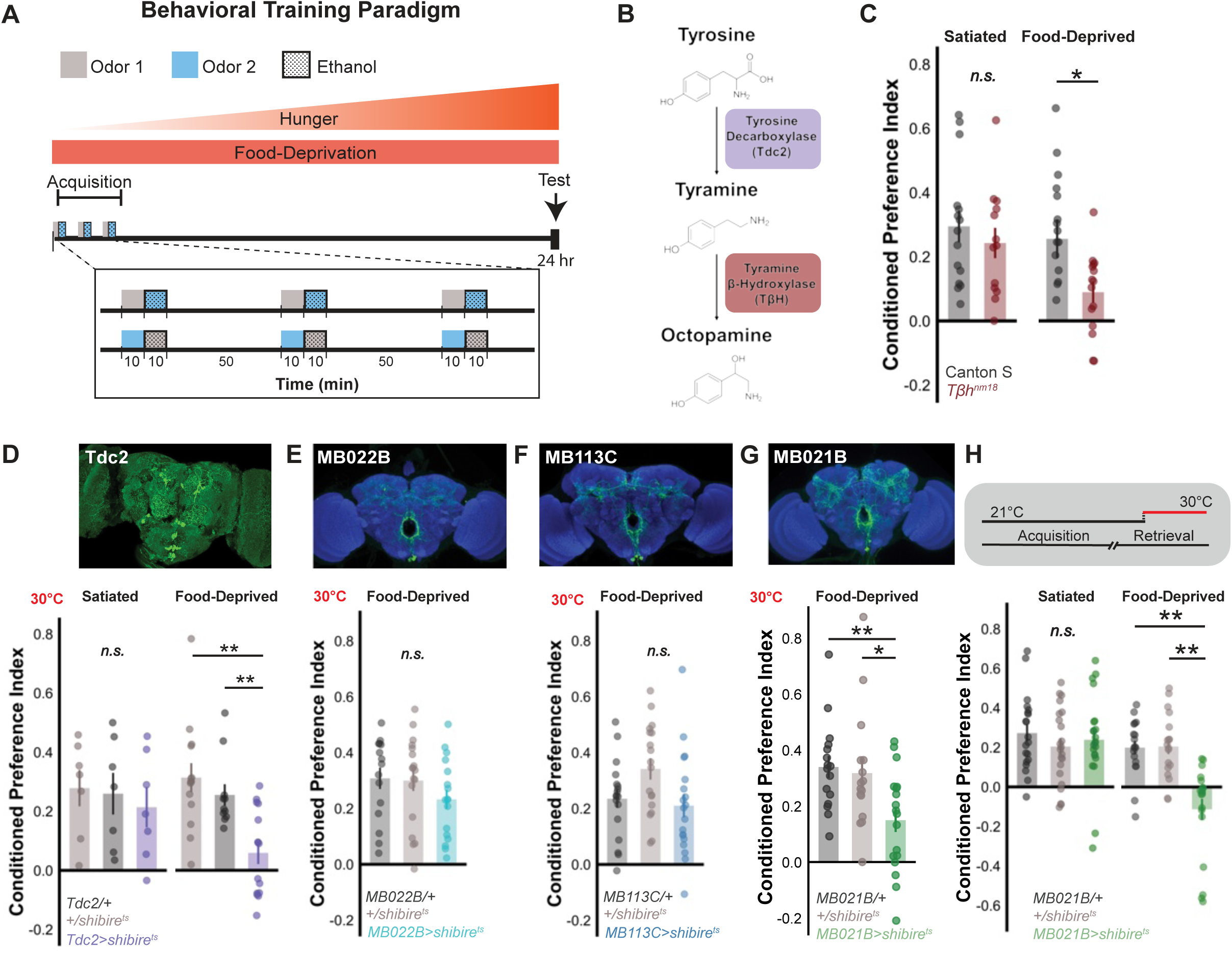
Octopaminergic neurons are necessary for food-deprived ethanol preference. (A) Schematic of ethanol conditioning paradigm used for food-deprivation experiments. Reciprocally trained vials of 30 flies are presented with three exposures of an unpaired odor and odor paired with an intoxicating dose of ethanol (66%, see methods) for 10 minutes each. There are 50-minute intervals of air exposures in between each pairings. ‘Food-deprived’ flies are placed on 2% agar vials prior to the start of training and kept on these vials until the test period, while ‘satiated’ flies are fed yeast pellets following the acquisition period. (B) Schematic of the octopa-mine/tyramine synthesis pathway detailing the conversion of tyrosine to tyramine with the tyrosine decarboxylase enzyme and subsequent conversion to octopamine via the tyramine β hydroxylase enzyme. (C) A tyramine β hydroxylase null mutant that is unable to convert tyramine to octopamine does not affect ‘satiated’ ethanol preference. However, this mutation exhibits a significant reduction in preference when flies are ‘food-deprived’ relative to Canton S wild-type controls. T(29)=2.3976, p<0.05. (D) Thermogenetic silencing of tyramine/octopamine expressing neurons throughout acquisition and test periods does not affect preference when the fly is ‘satiated’ but significantly reduces preference when the fly is ‘food-deprived’ relative to transgenic controls. One-way ANOVA with Tukey’s HSD method with Holm’s inference statistic was used to compare mean and variance across planned comparisons. F(2,38)=10.97, p<0.001. (E) Maximum intensity projection of MB113C-splitGAL4>20xUAS-IVS-CsChrimson-mVenus (green) and nc82 neuropil stain (blue) (images curated from Janelia Flylight Project Team; Aso et al., 2014a). Silencing MB113C-splitGAL4 using shibirets throughout acquisition and retrieval did not alter food-deprived ethanol preference relative to transgenic controls. (F) Maximum intensity projection of MB022B-splitGAL4>pJFRC200-10XUAS-IVSmyr::smGFP-HA (green) and nc82 neuropil stain (blue)(images curated from Janelia Flylight Project Team; Aso et al., 2014a). Silencing MB022B-splitGAL4 using shibirets throughout acquisition and retrieval did not result in food-deprived ethanol preference. (G) Maximum intensity projection of MB021B-splitGAL4>pJFRC200-10XUAS-IVSmyr::smGFP-HA (green) and nc82 neuropil stain (blue)(images curated from Janelia Flylight Project Team; Aso et al., 2014a). Silencing MB021B-splitGAL4 using shibirets throughout acquisition and retrieval did result in a significant food-deprived ethanol preference. One-way ANOVA with Tukey’s HSD method with Holm’s inference statistic was used to compare mean and variance across planned comparisons. F(2,49)=5.4586, p<0.01. (H) Thermogenetic inactivation of MB021B-splitGAL4 during the retrieval period results in a reduction of ethanol preference when compared to GAL4 controls only when flies are food-deprived. One-way ANOVA with Tukey’s HSD Method with Holm’s inference statistic was used to compare mean and variance across planned comparisons. F(2,54) = 17.8807, p < 0.001. Bar graphs illustrate mean +/-standard error of the mean. Raw data are overlaid on bar graphs. Each dot represents approximately 60 flies (30 per odor pairing). *p<0.05 **p<0.01 ***p<0.001.

Octopamine/tyramine expressing neurons comprise a large and heterogenous populations of neurons (Figure 1D). Due to this, we sought to define the smaller subset of octopamine/tyramine neurons that are mediating the observed state-dependent ethanol seeking. We investigated refined expression patterns of octopamine/tyramine expressing neurons (Figure 1D). Since the mushroom body and accompanying olfactory reward circuits play a pivotal role in ethanol preference memory^91,104,105^ we focused on *Tdc2*-expressing neurons that innervate mushroom body circuitry^69,73,83,98,106,107^. Neurons within the ventral paired median 3 (VPM3) and ventral paired median 4 (VPM4) clusters innervate dopaminergic neurons within the appetitive-encoding PAM and aversive-coding PPL1 clusters^73,83,106^ and are captured by the MB022B (VPM3/VPM4), MB021B (VPM4), and MB113C (VPM4) split-GAL4 driver lines^75^. Similar state-dependent effects have been mapped to the VPM4 neurons which directly modulate the γ1pedc>αβ mushroom body output neuron (MBON)^83^, sugar-sensing gustatory receptor neurons^85^, and appetitive DANs^73,106^. Thermogenetically inactivating neurons using the MB021B split-GAL4 (Figure 1G, Figure S2), but not the MB113C (Figure 1E) or MB022B (Figure 1F) split-GAL4 drivers reduced ethanol memory while the fly was food-deprived.

In our paradigm, flies are food deprived upon the start of training and hunger builds over the time until the testing (Figure 1A). If ethanol seeking resulting from inactivating MB021B is dependent on state, we would expect that the activity of MB021B neurons are required for ethanol memory retrieval and not acquisition. Indeed, inactivating MB021B during the odor choice 24 hours after training significantly reduced ethanol memory retrieval (Figure 1H), indicating that neurons within the MB021B expression pattern convey information that is necessary for ethanol preference only when the fly is food-deprived. Inactivating MB021B or MB113C neurons in satiated flies during training or test had no effect (Figure S2D,E).

### VUMa3 neuron activity is required for hunger-dependent ethanol memory

Our results demonstrated that inhibiting MB021B, but not MB113C neurons, reduced ethanol memory retrieval. This was surprising since both drivers were reported to have expression within the same VPM4 neurons^75^. To identify whether there were differences between the driver lines, we aligned their expression using Volumetric Visualization Device (VVD) Viewer and found that MB021B also has expression in an asymmetric VUMa3 neuron that is not present in MB113C (Figure 2A). This suggests that the observed state-dependent ethanol seeking phenotype is associated with the VUMa3 neuron. Unlike VPM4, VUMa3 has not been extensively studied beyond its initial characterization within the Tdc2-GAL4 expression pattern^69,107^.

**Figure 2.**
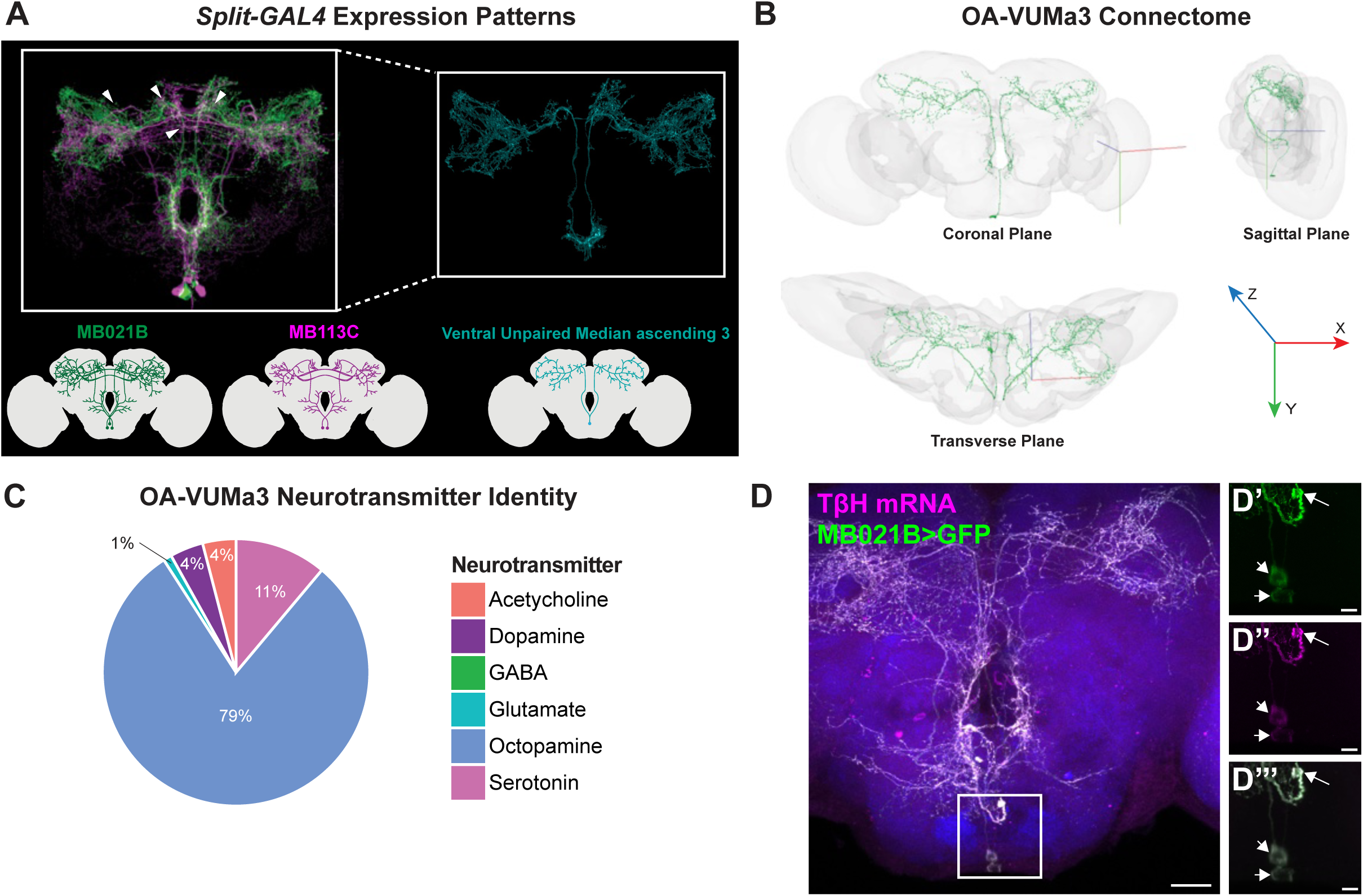
Octopamine-VUMa3 Neurons Release Octopamine for State-Dependent Autoregulation. (A) Schematic of neurons within the MB021B expression pattern (left) and neurons within the MB113C expression pattern (right). MB021B-splitGAL4>20xUAS-IVS-CsChrimson-mVenus (green) and MB113C-splitGAL4>20xUAS-IVS-CsChrimson-mVenus (magenta) expression patterns overlaid on top of each other using VVDViewer (Version 1.0.0). Maximum intensity projections of expression patterns are pseudo-colored from the FlyLight Project Team at Janelia Research Campus (Aso et al., 2014a). Extracted differences observed within the MB021B-spliGAL4 expression pattern that are not observed within MB113C-splitGAL4 are psuedo-colored in teal and characteristic features are annotated (white arrows). (B) Connectome reconstruction of OA-VUM neuron from FlyWire database from coronal, horizontal, and sagittal plane. (C) The pie chart depicts predicted neurotransmitter identity released from OA-VUMa3 synapses according to the FlyWire database. Octopamine is the predicted neurotransmitter at the majority (79%) of synapses, while serotonin comprises 11%, acetylcholine comprises 4%, dopamine comprises 4%, and glutamate comprises 1%. (D) Fluorescent in situ hybridization of MB021B-splitGAL4>10xUAS-myrGFP (green) using a fluorescent probe specific to tyramine β hydroxylase (magenta) indicates that 98Ma3 is octopaminergic. Neuropil stain (nc82) is shown in blue. The long arrow in D’-D’’’ shows 98Ma3, while short arrows show 9PM.4. Scale bars represent 20 µm (D) or 5 µm (D’-D’’’).

To gain a more detailed understanding of VUMa3 morphology, we took advantage of the *Drosophila* connectome and reconstruction tools available through the FlyWire database^108^. We focused on VUMa3 due to its morphological similarity with the identified VUMa3 neuron in the MB021B expression pattern (Figure 2A, 2B). Predictive neurotransmitter scores^173^ indicate that most of the output synapses from VUMa3 are octopaminergic (79%), with the second largest predicted neurotransmitter being serotonin (11%) (Figure 2C). Fluorescent *in situ* hybridization using probes specific for *Tβh*, the rate-limiting enzyme for octopamine synthesis^93,109,110^, revealed the VUMa3 neuron expresses the *Tβh* transcript (Figure 2D), thereby confirming VUMa3 expresses octopamine as a neurotransmitter.

### Octopamine in the VUMa3 neuron relays hunger-dependent ethanol memory

Our data suggest that VUMa3 conveys information about food-deprivation to influence expression of ethanol memory. We hypothesized that VUMa3 activity increases with hunger, and VUMa3 activity would build in the 24 hours between memory formation and expression.. To test this, we measured in vivo baseline calcium activity in the large VUMa3 axon tracts visible in the superior medial protocerebrum (SMP) following 24-hours of food-deprivation (Figure 3A). Since we were not evoking a response, baseline activity was quantified as a signal-to-noise ratio (SNR), in which GCaMP7b signal from VUMa3 was compared to the background signal of the brain (see Methods). A significant increase in SNR was observed in food-deprived flies compared to satiated flies (Figure 3B), suggesting that food-deprivation increases the baseline calcium activity of VUMa3 neurons across different laser powers.

**Figure 3.**
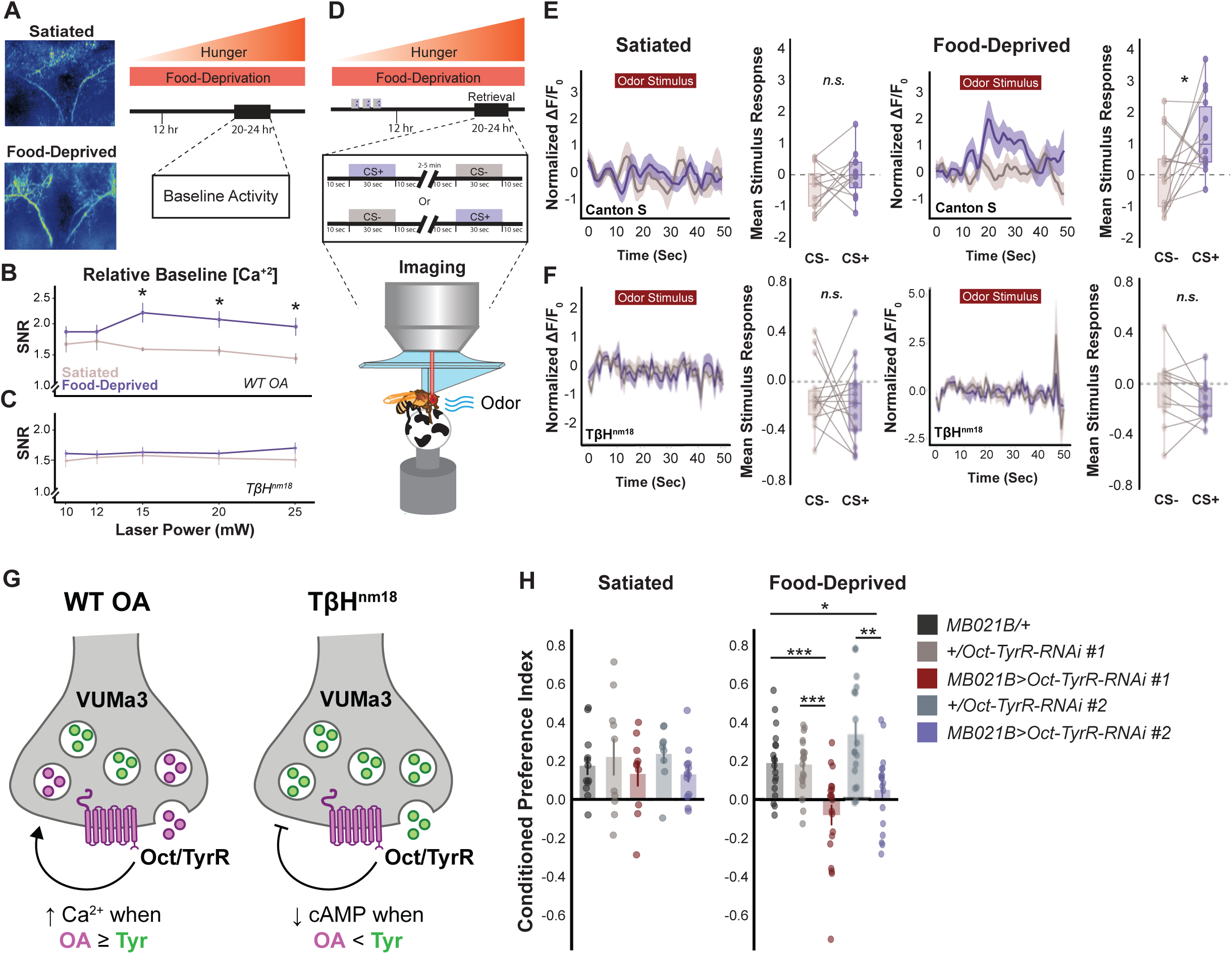
Octopamine-VUMa3 Neurons Exhibit State-Dependent Baseline Activity That Mediate Ethanol Memory Retrieval Through the Oct/Tyr Receptor. (A) Baseline signal-to-noise (SNR) measurements were taken of GCaMP7b-expressing flies that were either satiated or food-deprived for 20-24 hours at a range of laser powers: 10, 12, 15, 20, and 25 mW. Representative average projections of the calcium response to air baseline are displayed for satiated and food-deprived flies. (B) Food-deprived wild-type Canton S flies exposed to air across a range of laser powers show a significant increase in SNR ratio. Mixed models ANOVA with planned contrasts at 10, 12, 15, 20, and 25 mW indicate a significant interaction between condition and laser power F(4,48) = 2.8384, p < 0.05). All post hoc analyses were performed with Bonferroni corrections. Line plots show mean +/-standard error of the means. (n =7 flies within each condition, measured at each laser power). Significant differences were observed between food-deprived and satiated at 15 mW (p < 0.05), 20 mW (p < 0.05), and 25 mW (p < 0.05). (C) TβhnM18 flies did not show a significant change in SNR across food-deprived or satiated conditions. (D) Schematic of training paradigm used with MB021B-split-GAL4>UAS-GCaMP7b imaging experiments where flies are either satiated or food-deprived as in Figure 1. During the typical testing period (∼20-24 hours), flies are exposed to the paired odor (CS+) or unpaired odor (CS-) for 30 seconds under a 2-photon microscope. Flies were exposed to CS+ or CS-with the order of exposure being randomized, with a 2-5 minute rest interval in between. (E) Satiated flies exposed to CS+ or CS-odors show no strong calcium response to either odor. Average calcium traces of each response are overlaid on top of each other. Boxplots quantifying mean stimulus response across flies show that there was no significant difference in stimulus response of satiated flies between the CS– and CS+ odors. Food-de-prived flies show a strong time-delayed response to the CS+, and no response to the CS-. Results are quantified in the boxplots, with an onserved significant difference in stimulus response to the CS+ across food-deprived flies when compared to the CS-(T(13) = 2.78, p <0.05). (F) Satiated or food-deprived TβhnM18 flies exposed to CS+ or CS-odors show no strong calcium response to either odor. Average calcium traces of each response are overlaid on top of each other. Boxplots display the mean stimulus response across flies to either the CS– and CS+ odors. (G) Schematic of model displaying the autoregulation of VUMa3(mx) neurons during wild-type and octopamine-null conditions, where autocrine octopamine release increases baseline activity through the Oct/Tyr Receptor. (H) Oct/TyrR RNAi-mediated knockdown in VUMa3 results in ethanol preference when flies are satiated, but results in a significant decrease in ethanol preference relative to transgenic controls. The food-deprived condition was analyzed using one-way ANOVA with Wilcoxon/Kruskal-Wallis tests to compare mean and variance across groups. H(4,93) = 27.7228, p<0.0001. Bar graphs illustrate mean +/-standard error of the mean. Raw data are overlaid on bar graphs. Each dot represents approximately 60 flies (30 per odor pairing). *p<0.05 **p<0.01 ***p<0.001.

These data suggest that VUMa3 becomes recruited into ethanol memory circuitry as hunger builds. As expected, the VUMa3 neuron did not show significant differences in activity to the unpaired (CS-) or paired (CS+) odor cue in satiated flies (Figure 3D-E), whereas it showed significantly higher activity in response to the CS+ compared to the CS-in food-deprived flies (Figure 3E). Thus, VUMa3 conveys information about the CS+ that is necessary for ethanol preference only when the fly is hungry.

To examine the role of octopamine/tyramine signaling from VUMa3 in conveying state-dependent information, we measured baseline calcium activity in fed and food-deprived Tβh^nM18^ flies. Contrary to flies exhibiting functional levels of octopamine, we did not observe a significant increase in baseline activity in VUMa3 starved Tβh^nM18^ flies relative to the fed state (Figure 3C). Similarly, neither fed nor starved Tβh^nM18^ flies showed significant differences in VUMa3 activity when exposed to a CS+ and CS-(Figure 3F), indicating that the state-specific information conveyed by VUMa3 is dependent on octopamine signaling. This result indicates that Tβh function is required for VUMa3 to distinguish between a fed or starved internal state. We hypothesize that this starvation-induced increase in VUMa3 activity is likely due to autocrine binding of octopamine released from VUMa3 itself due to the very low number of presynaptic octopaminergic inputs onto VUMa3 (Figure S3).

Since Tβh^nM18^ flies lack octopamine and show an eightfold increase in tyramine^93^, our results could be interpreted either as a consequence of decreased octopaminergic effects or increased tyraminergic effects on VUMa3. To identify whether octopamine or tyramine primarily mediates starvation-dependent responses in VUMa3, we sought to identify potential octopamine or tyramine receptors that regulate VUMa3 signaling in an autoregulatory fashion, and in a state-dependent manner.

The Octopamine-Tyramine Receptor (Oct-TyrR, *honoka*) is receptor that responds to both octopamine and tyramine. It is structurally and pharmacologically analogous to alpha-2 adrenergic receptors^112,113^, and acts as a positive regulator upon octopamine binding, and a negative regulator upon tyramine binding^112,114,119–125^. It has also been found to alter motor neuron activity in response to nutritional state ^111^. We reasoned that Oct-TyrR could help distinguish whether the increase we observe in VUMa3 activity upon food deprivation is due to an increase in octopamine-mediated activation or a decrease in tyramine-mediated attenuation (Figure 3G).

Consequently, upon Oct-TyrR knockdown, we expected either a decrease in ethanol-seeking if the starvation-dependent effects were primarily mediated by octopamine, or an increase/no change in ethanol seeking if the starvation-dependent effects were primarily due to tyramine. Importantly, knockdown of Oct-TyrR in VUMa3 neurons using MB021B resulted in a significant decrease in state-dependent ethanol seeking in food-deprived flies (Figure 3H), suggesting that octopamine primarily acts via autoregulation through Oct-TyrR to modulate starvation-dependent effects on ethanol seeking behaviors in our VUMa3 circuit. While unlikely based on previous experiments, it is important to note that VPM4 neurons in MB021B also express octopamine and could be contributing to the observed Oct-TyrR-mediated effect rather than simply VUMa3 autocrine signaling.

### Post-synaptic partners of VPM4 and VUMa3 neurons

To identify downstream partners of VUMa3, we compared post-synaptic partners of MB021B and MB113C neurons using the transsynaptic labeling tool *trans-* Tango^127,128^ (Figure 4A). Both MB113C and MB021B innervated regions that are associated with olfactory memory (Figure S4) including the calyx (Figure S4), alpha/beta (α/β), alpha’/beta’ (α’/β’), gamma (γ) lobes of the mushroom body (Figure S4), and the antennal lobe (AL) (Figure S4). Additionally, MB113C and MB021B innervate similar populations of neurons within the superior lateral protocerebrum (SLP), anterior ventrolateral protocerebrum (AVLP), lateral horn (LH), posterior lateral protocerebrum (PLP), saddle (SAD), pars intercerebralis (PI), and peduncle (PED) (Figure S4). These similarities across the two populations can be attributed to VPM4 neurons and illustrate the broad modulatory control octopaminergic neurons have across the brain.

**Figure 4.**
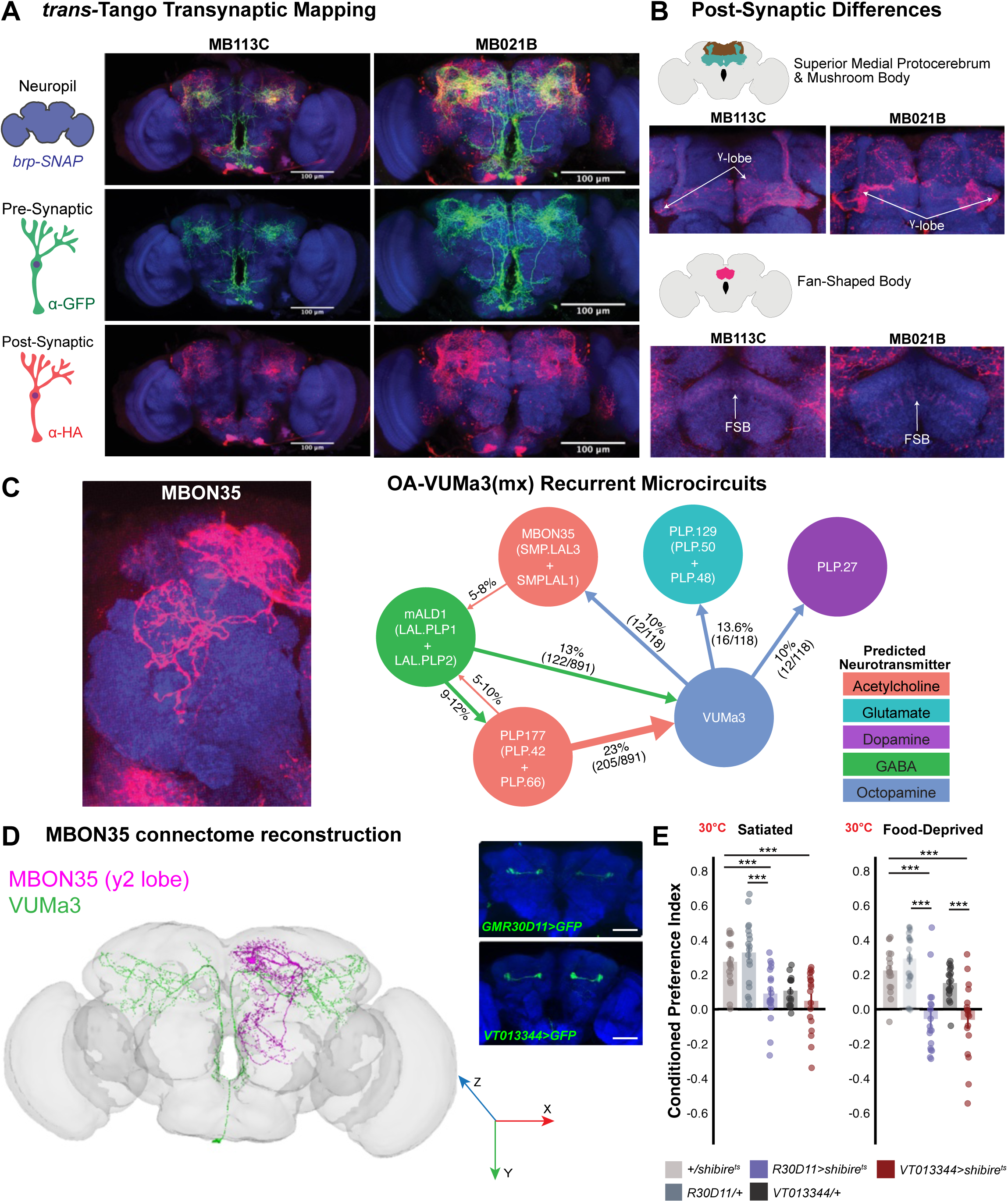
Post-synaptic targets of MB113C and MB021B. (A) Using the trans-Tango transsynaptic labeling tool allows for the visualization of pre-synaptic neurons (green), their post-synaptic targets (red), and neuropil (blue). Representative maximum intensity projections of MB113C and MB021B and their subsequent separate channels are displayed. (B) Differences in post-synaptic targets between MB113C and MB021B are observed across three specific brain regions. Specifically focusing on the superior medial protocerebrum (SMP) and Mushroom Body (MB) lobes, we observed that MB021B consistently had comparatively stronger innervation of the y-lobe and particularly in the dorsal region and near the peduncle (PED). Within the fan-shaped body (FSB) we observed that MB113C innervated layers 5-6, while MB021B innervated layer 4. These differences could be attributed to VUMa3 rather than VPM4. (C) MB021B displayed MBON35 as a putative post-synaptic target that isn’t in MB113C post-synaptic targets (shown in red). Schematic of a microcircuit comprising two of the top inputs onto VUMa3 (shown as a percentage of total inputs onto VUMa3), along with the top 3 outputs from VUMa3 containing >10 synapses (shown as a percentage of total outputs from VUMa3). Each cell node is pseudo-colored with predicted neurotransmitter type, and reciprocal connections between these populations are displayed. Insets on the right represent potential examples of the top outputs shown within the schematic from trans-Tango labeling experiments using MB021B (postsynaptic targets in red and neuropil in blue). (D) Coronal display of MBON35 connectome reconstruction from FlyWire database. R30D11-GAL4 and VT013344-GAL4 were previously shown to contain MBON35. Maximum intensity projection of R30D11-GAL4>10xUAS-myr::GFP and VT013344-GAL4>10xUAS-myr::GFP (green) and nc82 neuropil stain (blue). Scale bars represent 50 µm. (E) Silencing of R30D11-GAL4 and VT013344-GAL4 using shibirets resulted in significant reduction for ethanol preference in both satiated (F(4,85) = 17.2574, p<0.0001) and food-deprived (F(4,84) = 11.7569, p<0.0001) flies. The food-deprived condition had a larger reduction relative to transgenic controls. Bar graphs illustrate mean +/-standard error of the mean. Raw data are overlaid on bar graphs. Each dot represents approximately 60 flies (30 per odor pairing). *p<0.05 **p<0.01 ***p<0.001.

It is worth noting that some differences in MB113C and MB021B targets also emerged: MB021B exhibits stronger expression in the dorsal γ-lobe, γ1-peduncle, γ2 compartments (Figure 4B) and increased superior medial protocerebrum (SMP) innervation (Figure 4B). Importantly, while both drivers also synapse onto neurons with arborizations in the fan-shaped body (FSB), they seem to target different layers or populations of neurons (Figure 4B). Specifically, VUMa3 seems to synapse onto a neuron that has compartmentalized arborization patterns within layer 5 of the FSB.

Finally, the most visible difference between the post-synaptic partners of MB113C and VUMa3 is the density of innervation surrounding the pedunculus observed in the post-synaptic partners of VUMa3 (Figure 4A, S4). Specifically, VUMa3 synapses onto several neurons that have arborizations in the superior and inferior clamp regions.

These data demonstrate that VUMa3 has a vast arborization pattern and innervates divergent brain regions across several sensory modalities. This includes intrinsic and extrinsic neurons within the γ-lobe of the mushroom body and predominantly tangential layer 4 of the FSB. The differences in the FSB layers innervated by MB113C and VUMa3 may highlight the importance of state-dependent modulation within this convergent region^128^, and could act to bias directional/motor behaviors towards an essential resource given the state of the animal^129^. These tangential FSB layers tend to be downstream of the MB and LH circuitry and are vital for olfactory navigation^130^, foraging^131^, and innate and conditioned avoidance behaviors^132^. Thus, state-dependent VUMa3 modulation in these downstream circuits may be another manner of fine-tuning behavioral output.

We confirmed the post-synaptic connections of VUMa3 revealed by *trans-*Tango in the FlyWire connectome, and identified pre-synaptic targets to identify recurrent networks^106,108,133,134^. The top synaptic input onto VUMa3 is from PLP177, a cholinergic neuron with extensive arborizations in the PLP^133^ (neuprint.janelia.org^135^; Figure 4C). The second-largest input onto VUMa3 is from the GABAergic neuron mALD1(CREa1-ventral). These neurons form a microcircuit with each other and MBON35, one of the top targets of synaptic outputs of VUMa3 (Figure 4C).

MBON35 (Figure 4C-D) is an ‘atypical MBON’ that compartmentalizes to the γ2 compartment of the mushroom body but also has dendritic arbors outside of the mushroom body^106^. We sought to determine whether silencing MBON35 could recapitulate the state-dependent ethanol seeking observed when VUMa3 is silenced by using two GAL4 lines identified using NeuronBridge^136^ that drive expression in MBON35 (R30D11 and VT013344). Thermogenetic inactivation of MBON35 throughout our conditioning and retrieval paradigm displayed reductions in ethanol seeking in both satiated and hungry flies (Figure 4E). However, blocking MBON35 reduced ethanol seeking more severely in hungry than sated flies, suggesting that MBON35 may be partially responsible for modulating state-dependent behaviors. Due to MBON35 being a primary synaptic output of the VUMa3 neuron and its displayed role in ethanol seeking behavior across ‘hunger states,’ we suggest that VUMa3’s state-dependent modulation is upstream of MBON35 and primarily responsible for our observed behavioral effects and differences across hunger-states.

### VUMa3 State-Dependent Circuit

The VUMa3 state-dependent circuit described in this study highlights a parallel ethanol memory circuit motif that is only active upon food-deprivation to drive ethanol seeking in *Drosophila.* In this study, we observe that VUMa3 baseline activity correlates with increasing hunger and has a role in state-dependent modulation of memory circuits. One can postulate that state-dependent modulation of this manner can have maladaptive effects and affect ethanol memory and seeking. Along those lines, noradrenergic signaling has been previously shown to be upregulated during alcohol intoxication^137,138^ and is an enticing candidate for understanding how internal state influences the development of dependency^41,139^.

The vast arborization, morphology, and dense innervation observed with VUMa3 neurons is a characteristic reminiscent of noradrenergic neurons in other organisms, and spans several structures required for memory and sensory processing^42–48,154–158^. We speculate that the VUMa3 state-dependent circuit is not solely integrating hunger state to drive ethanol seeking but can be expanded to other internal states and types of olfactory memories. Our study provides a potential circuit motif that directly links internal state-modulation to memory processing, as well as an understanding of the impact these state-dependent circuits have in mediating drug use, addiction, cravings, and relapse. Thus, providing a potential mechanism by which stress-arousal systems affect memory processing and retrieval mechanisms, and how these innately beneficial systems may be co-opted to result in maladaptive behaviors, cravings, and relapse.

Further characterization of this circuit provides an opportunity to assess how responsive the VUMa3 circuit is to other stressors, how it influences other behaviors, and if this circuit can be co-opted for stress-induced reinstatement of ethanol preference. Not all stressors might function to modulate behaviors similarly, but internal states that have competing drives may directly oppose each other to influence behavior within the same circuit. Sophisticated experiments furthering our understanding of this circuit, as well as other circuits exhibiting similar state-dependencies, can potentially be used to initiate stress-induced alcohol seeking and find means to counter these effects.

## Lead contact

Further information and requests for resources and reagents should be directed to and will be fulfilled by the lead contact, Karla R. Kaun.

## Materials availability

This study did not generate new unique materials or reagents.

## Data and code availability

Raw and analyzed data for all figures are openly accessible and available through a Brown Library Data Management link at time of full publication. Source data files for visualization and analysis of data are openly accessible and available at https://github.com/kaunlab.

## Acknowledgements

This work was funded by the National Institute on Alcohol Abuse and Alcoholism (R01 AA024434 to K.R.K. and F99 NS118741 to K.M.N.), the National Institute of General Medical Sciences (R01 GM115510 to K.R.K.), the National Institute on Aging (T32 5T32AG041688 supporting L.M.S.), the National Institute on Deafness and Other Communication Disorders (R01 DC017146 and R01 DC020703 to G.B), the Hubert & Richard Hanlon Trust and Howard Hughes Medical Institute (Gilliam award to K.M.N.), and an Innovation Award from the Carney Institute for Brain Science at Brown University (to G.B.).

We would like to thank Dr. Carolina Haas-Koffler (Brown University), Tara White (Brown University), and John McGeary (Brown University) for their insight on the translational impact of how stress impacts noradrenergic signaling in the context of alcohol use disorder. We would also like to thank Dr. Sarah Certel (University of Montana) and Dr. Steve Stowers (Montana State University) for their insight into the morphology and function of octopamine neurons in the *Drosophila* brain. Thanks to Dr. Mareike Selcho (Leipzig University) for insight on octopamine neuron morphology and connectomics. Thanks to Dr. Rebecca Oramas for assistance with figure design and arrangement. Thanks to all members of the Kaun Lab who provided helpful feedback on all aspects of the science performed in association with this work.

## Author contributions

K.M.N. conceptualized and designed experiments, performed data acquisition, analysis, and visualization, wrote, and revised the manuscript, and provided mentoring.

L.M.S. conceptualized and designed experiments, performed data acquisition, analysis, and visualization, wrote, and revised the manuscript, and provided mentoring.

A.W. performed data acquisition under the mentorship of K.M.N. and K.R.K.

S.S. performed data acquisition under the mentorship of K.R.K.

V.M.C. performed data acquisition under the mentorship of L.M.S. and K.R.K.

M.T provided *Tango* reagents, advice, and feedback on the manuscript.

G.B. provided *Tango* reagents, advice, and feedback on the manuscript.

K.R.K. conceptualized and designed the research study, performed data acquisition and analysis, provided advice on data analysis and visualization, wrote and revised the manuscript, provided funding to perform the study, and provided mentoring.

## Material and methods

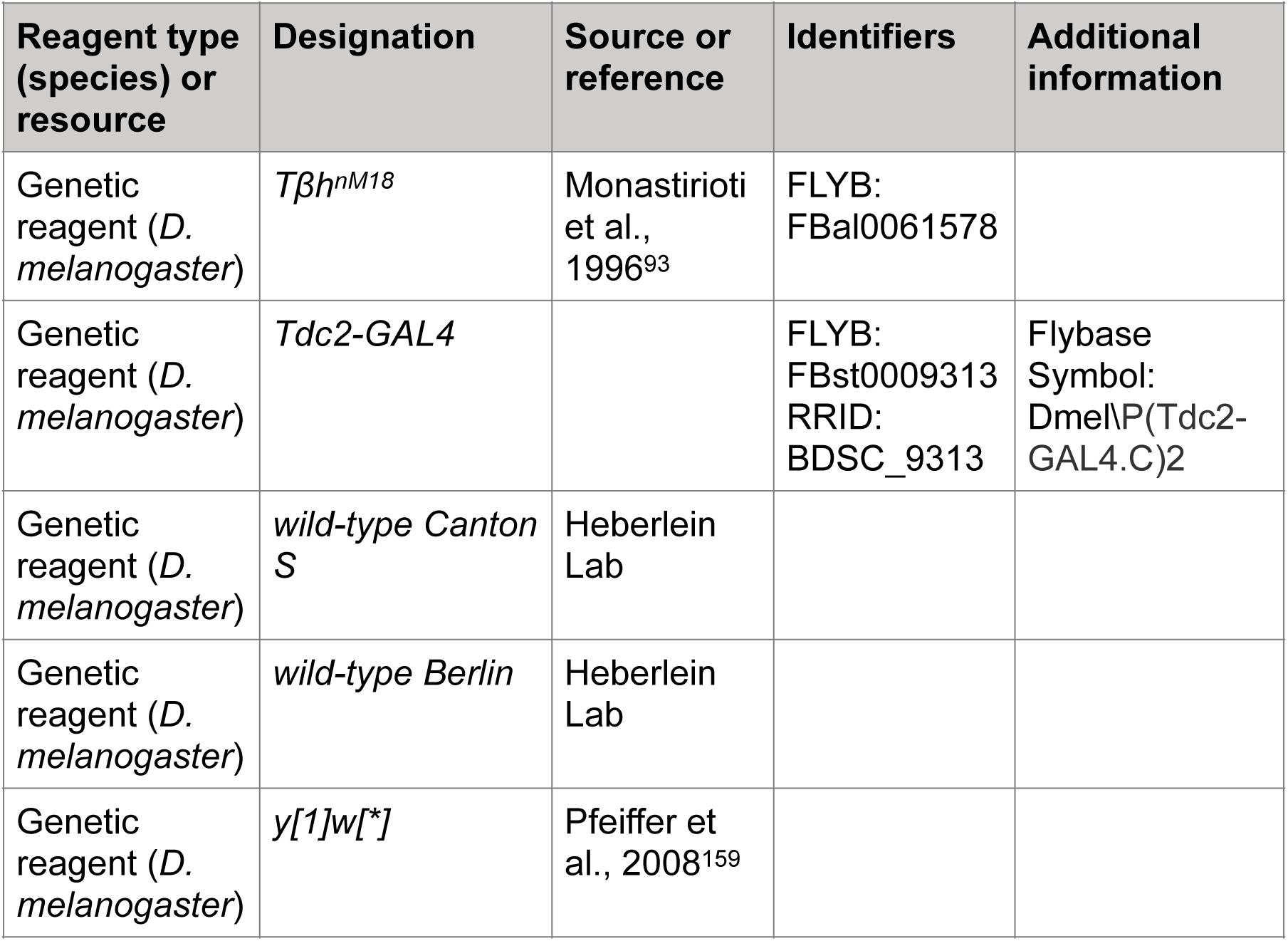
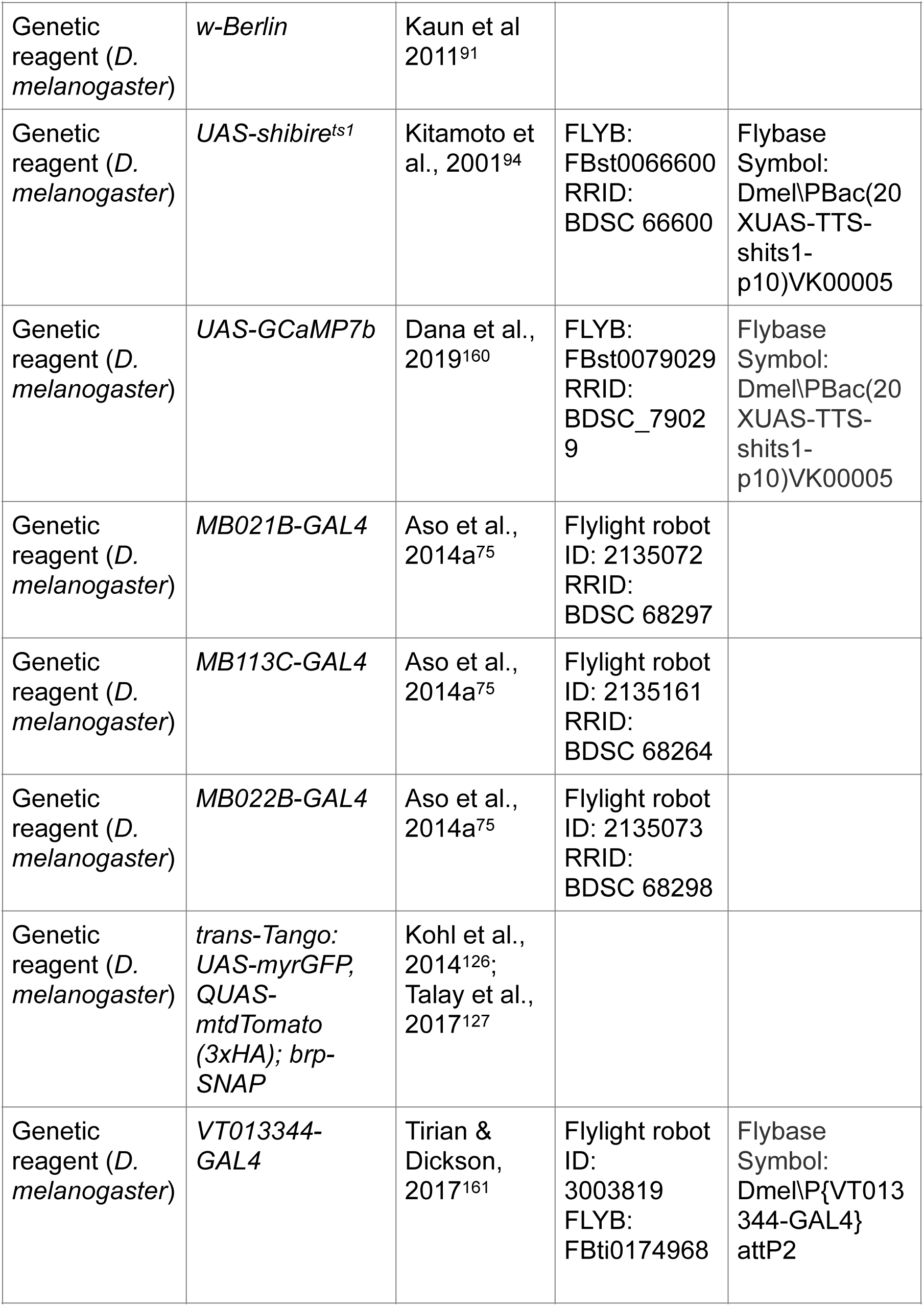
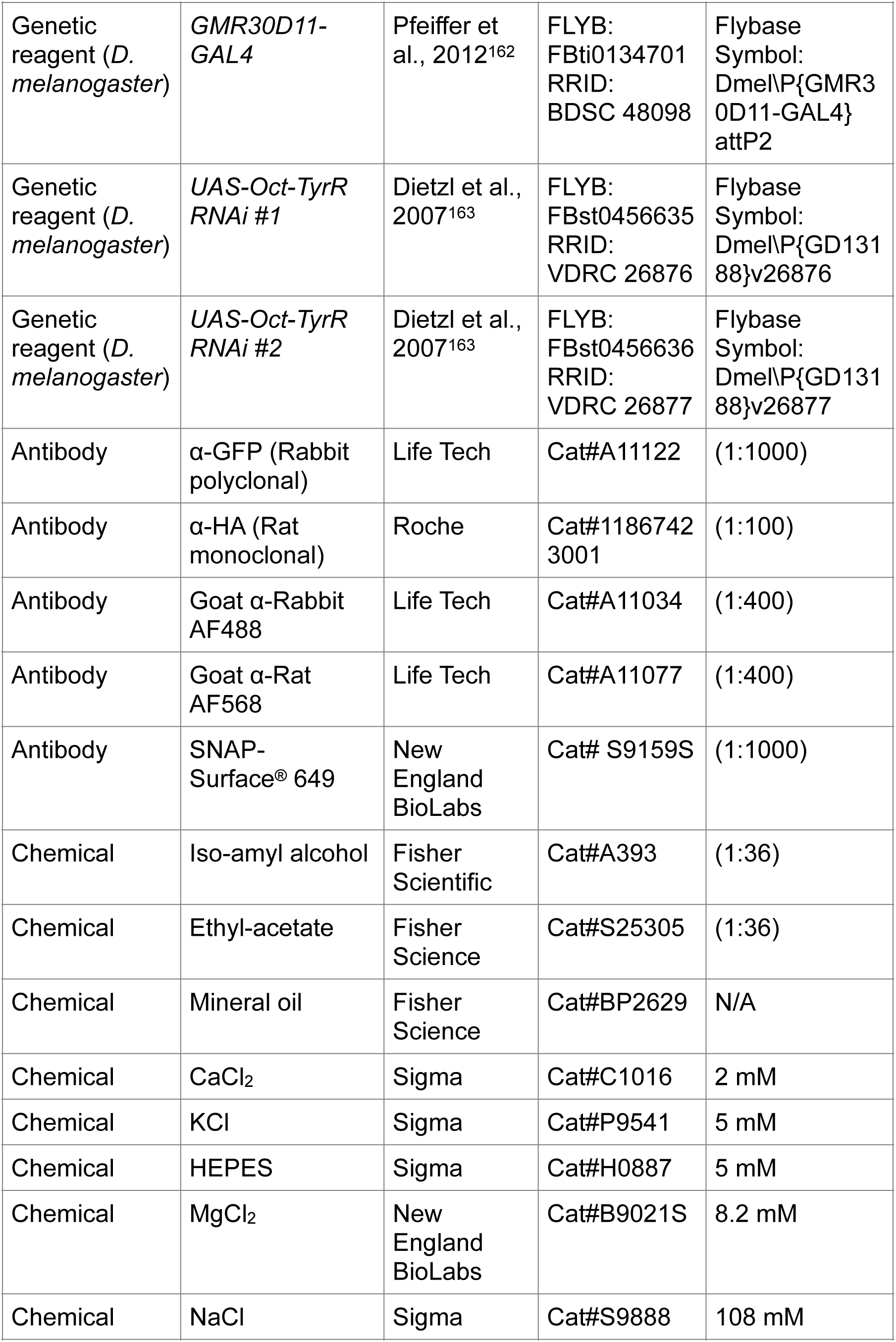
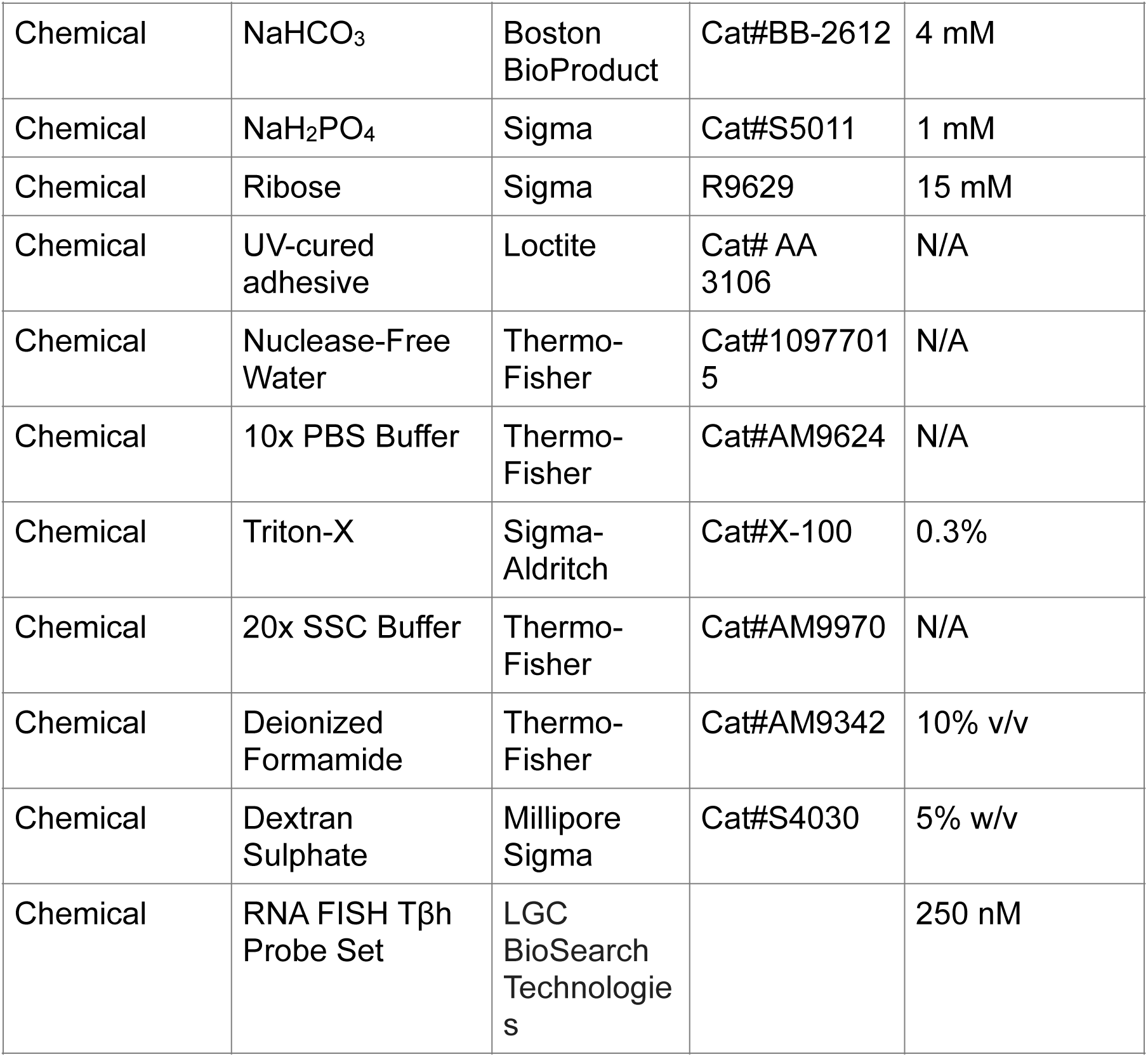
Key Resources table.

### Fly Stocks and Conditions

All *Drosophila melanogaster* lines were raised on standard cornmeal-agar media with tegosept anti-fungal agent and maintained at either 18°C, 24°C, or 30°C in 14:10 hour light:dark cycles. Complete list of fly stocks used in the study are listed in the Key Resources Table.

### Thermogenetic Inactivation

Thermogenetic inactivation experiments required maintaining flies at 30°C post-conditioning or prior to testing odor-cue preference, and instances of this are stated as such in the figure legends and accompanying results. For the 30°C post-conditioning, temperatures were maintained in humidity-controlled chambers under 14:10 hour light:dark cycles.

### Food-Deprivation

To assess the state-dependent role modulatory neurons have in ethanol memory we modified our lab’s previously established paradigm^91,92,105^. Specifically, we sought to parse out the impact different food-deprivation paradigms have on ethanol memory. Flies considered to be in the ‘satiated’ state are trained in perforated culture vials containing 1% agar and supplemented with yeast post-training, similar to the established paradigm in our lab^91,92,105^. Flies were considered to undergo ‘food-deprivation’ when they were not supplemented with yeast post-training, essentially being food-deprived throughout acquisition and retrieval periods (Figure 1A).

All *Drosophila melanogaster* males undergoing food-deprivation were wet-starved in 14 ml culture vials containing perforated mesh lids, the same vials used for odor preference conditioning. These vials contained 1 ml of 1% agar solutions, same as used for training. Under standard satiated conditions, and during the ‘early’ food-deprivation paradigm, flies were supplemented with yeast pellets post-training while on 1% agar (Figure S1). Flies undergoing the 48 hour or ‘late’ food-deprivation paradigm were not supplemented with yeast pellets post-training and thus food-deprived throughout acquisition and retrieval periods, with the 48 hour paradigm adding an additional 24 hour food-deprivation period prior to the start of conditioning (Figure S1). For all subsequent experiments, the ‘late’ food-deprivation protocol was used (Figure 1A, S1).

### Odor Preference Conditioning

All *Drosophila melanogaster* lines, crosses, and progeny used for behavioral experiments were male flies collected 1-2 days post-eclosion and shifted to 18°C, 65% humidity, 14:10 hour light:dark cycle controlled chambers. Odor conditioning experiments were performed as described in Kaun et al, 2011^91^ and Scaplen et al, 2020^105^. Briefly, groups of 30 flies were placed on 1% agar in perforated 14 mL culture vials containing mesh lids to allow for proper exposure to odorants, humidified air, and/or volatilized ethanol. Behavioral training chambers were maintained at 65% humidity, under red-light illumination, and constant temperature (either at 21°C or 30°C, dependent on thermogenetic perturbation). Training chambers are described in more detail in Nunez et al., 2018^92^, but in short, vials of flies are conditioned within Plexiglass chambers (30 × 15×15 cm). Prior to training, vials are habituated in the training chambers for 20 minutes with humidified air (flow rate of 130 U). A training session consists of 10 minutes of an odor stimulus (flow rate of 130 U), followed by presentation of a different odor stimulus (flow rate: 130 U) paired with volatilized ethanol (flow rate of 90 U:60 U, ethanol:air). There is then a subsequent 50 minute rest period, where the chambers receive humidified air (flow rate of 130 U). Training sessions were repeated two more times for a total of three spaced training sessions. Fly vials in a separate training chamber are simultaneously receiving reciprocal odor pairings/training. Reciprocal training allows us to compare training chambers and account for any differences in innate odor preference. Odors used for experiments are 1:36 (odor:mineral oil) mixtures with ethyl acetate or isoamyl alcohol. Reciprocally trained odor vials were tested 24 hours later in the vertical Y maze for odor preference by streaming either odor (flow rate of 10 U) into the separate arms of the Y maze containing a collection vial. Flies climb into the collection vials at either arm of the Y maze, indicating their odor choice. A preference index is then calculated by counting the: (# of flies in the paired odor vial – # of flies in the unpaired odor vial)/total # of flies that climbed into the collection vials. A conditioned preference index (CPI) is calculated by averaging across the reciprocally trained groups.

### Odor Reactivity

Odor reactivity was evaluated for lines that exhibited an effect to thermogenetic inactivation and/or food-deprivation conditions and thus tested at the restrictive (30°C) or permissive (21°C) temperatures and/or food-deprived as described in odor preference conditioning section. Odors used were 1:36 (odor:mineral oil) mixtures of either isoamyl alcohol or ethyl acetate. Groups of 30 naïve male flies were presented with either an odor or humidified air at separate arms of the Y maze (flow rate of 10). A preference index was calculated by counting: (# flies in odor vial – # flies in the air vial)/total # flies that climbed into collection vials. These experiments were used to determine if either experimental perturbation disturbed odor perception or sensitivity.

### Fluorescence In Situ Hybridization (FISH)

Whole adult brain FISH was performed as previously described (Yang et al, 2017^109^). 5–10-day-old adult *Drosophila* male brains were dissected and fixed in 4% paraformaldehyde (Electron Microscopy Sciences #15710) for 20 minutes at room temperature. Brains were then washed with nuclease-free 1x PBS solution (Thermo-Fisher #10977015; Thermo-Fisher #AM9624), followed by a 40-minute wash in nuclease-free 1x PBS solution with 0.3% Triton-X (Sigma-Aldrich #X-100) to permeabilize the tissue. Samples were incubated in wash buffer (2x nuclease-free SSC solution (Thermo-Fisher #10977015; Thermo-Fisher #AM9970) and 10% v/v deionized formamide (Thermo-Fisher #AM9342)) for 10 minutes at 37°C. Wash buffer was subsequently removed and replaced with hybridization buffer (2x nuclease-free SSC solution with 10% v/v deionized formamide, 5% w/v dextran sulphate (Millipore Sigma #S4030), 250 nM FISH probes (LGC BioSearch Technologies) and primary antibodies (monoclonal rabbit anti-GFP (1:350, Life Technologies #A-11122), mouse anti-bruchpilot (1:40, mAb nc82, Developmental Studies Hybridoma Bank developed under the auspices of the NICHD and maintained by the Department of Biology, University of Iowa (Iowa City, IA))), and incubated overnight at 37°C with gentle nutation. Hybridization buffer was removed after 24 hours and samples were subsequently washed 3x for 10 minutes in wash buffer and incubated for 40 minutes in wash buffer containing secondary antibodies (AlexaFluor Goat anti-Rabbit 488 (1:200, Thermo-Fisher #A-11008) AlexaFluor Donkey anti-mouse 647 (1:200, Thermo-Fisher #A-21202)). Samples were subsequently washed 3x in wash buffer, mounted on a poly-L-lysine coated coverslip (Sigma-Aldrich #P1524; Corning #2845-22) in Fluoromount-G mounting medium (SouthernBiotech #0100-01) and imaged.

### FISH probe Sequences

Oligonucleotide probe set for TβH was identical to that used in Meissner et al, 2019^110^. The FISH probe set was purchased from LGC BioSearch Technologies (Novato, CA, USA) prelabeled with Quasar-570 dye. Sequences for the FISH probe set are provided in Supplementary file 1.

### Trans-Tango Immunohistochemistry

Experiments were performed according to the published FlyLight protocols with minor modifications that are described thoroughly in Scaplen et al, 2020^128^. All incubations/washes were done with the samples nutating to allow adequate exposure to molecular compounds. All *Drosophila melanogaster* lines, crosses, and progeny used for *trans-*Tango experiments were raised and maintained at 18°C in humidity-controlled chambers under 14:10 hour light:dark cycles on standard cornmeal-agar media with tegosept anti-fungal agent.

Briefly, adult flies that are 7-14 days old were anesthetized on ice, de-waxed in 70% ethanol and dissected in cold Schneider’s Insect Medium (S2). After dissection, tissues were incubated at 4°C for 22-26 hours in 1.2% cold paraformaldehyde (PFA) diluted in S2 solution. After fixation, samples were rinsed with cold phosphate buffered saline with 0.5% Triton X-100 (PBT) and subsequently washed 4 times for 15 minutes at 4°C. Following PBT washes, PBT was removed and samples were incubated in SNAP substrate diluted in PBT (SNAP-Surface649, NEB S9159S; 1:1000) for 1 hour at room temperature. Samples were then rinsed and washed 3 times for 10 minutes at room temperature and subsequently blocked in 5% heat-inactivated goat serum in PBT for 90 minutes at room temperature and incubated with primary antibodies (Rabbit α-GFP Polyclonal (1:1000), Life Tech #A11122, Rat α-HA Monoclonal (1:100), Roche #11867423001) for 48-72 hours at 4°C. Following removal of primary antibodies, samples were then rinsed and washed 4 times for 15 minutes in 0.5% PBT and incubated in secondary antibodies (Goat α-Rabbit AF488 (1:400), Life Tech #A11034, Goat α-Rat AF568 (1:400), Life Tech #A11077) diluted in 5% goat serum in PBT for 48-72 hours at 4°C. Samples were then rinsed and washed 4 times for 15 minutes in 0.5% PBT at room temperature and prepared for mounting in Fluoromount-G (SouthernBiotech, #0100-01) on poly-L-lysine-coated coverslips (Sigma-Aldrich #P1524; Fisher Scientific #12-541, square, 18mm, no. 1.5).

Confocal images were obtained using the Olympus FV3000 confocal microscope in Brown University’s Leduc Bioimaging Facility. Images were captured with 60x objective at 800 x 800-pixel resolution, with incremental laser power adjustments (Bright-Z) that occur with progress through the z-plane. The incremental adjustments in laser power with z-depth aided generation of a homogeneous signal where it seemed necessary. Stacks of 0.48-1.00 µm z-slices were acquired with optimal isotropic resolution in the x-y planes and were subsequently stitched together. All maximum z-projection stacks used in figures were completed using FIJI (http://fiji.sc).

To identify differences in expression patterns between MB021B and MB113C split-GAL4s, we used an open-source adaptation of Fluorender^164^ (VVDViewer (https://github.com/takashi310/VVD_Viewer/releases) to segment the arborization patterns of the split-GAL4 patterns (Aso et al., 2014^75^; Janelia Flylight Project Team). Brains were aligned to the JFRC 2018 template brain for comparison across samples^165,166^ (Figure 2A).

### Calcium Imaging

For imaging experiments, UAS-GCaMP7b were crossed with MB021B-GAL4. Male flies were collected 2-3 days post-eclosion and imaged at 5-10 days of age. Calcium imaging experiments were done similarly to those described in Scaplen et al, 2020^105^ with a few modifications.

Briefly, flies were anesthetized on ice and mounted onto an experimental holder (Weir et al., 2016^167^) with UV-activated Loctite light cure adhesive (Loctite AA 3106) using brief 3-6 sec pulses of UV light. Heads were tilted at a slight angle to open an imaging window in the posterior portion of the fly head. All legs, antenna, and the proboscis were left intact and unglued to not disturb flies. Once a viewing window was cut open in the cuticle using sharpened #5 forceps, cold Ribose-Substituted *Drosophila* Adult Hemolymph-Like Saline (AHLS) solution was added to the brain. *Drosophila* AHLS solution was modified slightly from standard CSHL protocol^168^, where 10mM sucrose and 5mM trehalose were substituted with 15mM ribose (in line with other starvation calcium imaging protocols^85^). The fly preparation was positioned on a customized stand underneath the two-photon scope (http://ptweir.github.io/flyHolder/^167^). The position of the ball and the stream delivery tubes were manually adjusted to the fly’s position in the holder.

A singular odor or volatilized ethanol stimuli were delivered within a 2 cm distance from the fly antenna (flow rate of 5:5, odor:air). Exposures to odor stimuli that are paired with volatilized ethanol were also delivered at a 2 cm distance from the fly antenna (flow rate of 5:5, odor:ethanol). Odors used were 1:36 (odor:mineral oil) mixtures with ethyl acetate or iso-amyl alcohol. Calcium imaging recordings were performed with a two-photon resonance microscope (Scientifica). Fluorescence was recorded from the neurons in the MB021B-GAL4 expression pattern (VPM4 and VUMa3), focusing on the neuropil and large medial axons from VUMa3 neurons in the superior medial protocerebrum of the brain.

All calcium imaging recordings were acquired using SciScan Resonance Software (Scientifica). The 2-photon laser had a wavelength of 930 nm at an intensity of 10-25 mW. Images were acquired at 512 x 512-pixel resolution with an average of ∼30.9 frames per second across 26(z) slices for a total 1.19 Hz using a Piezo to conduct volumetric imaging. Retrieval experiments were conducted across ∼50 sec (total # of cycles, 60): ∼10 sec (12 cycles) of air, 30 sec of odor stimulus (36 cycles), ∼10 sec (12 cycles) of air. Additionally, for retrieval experiments all flies were exposed to both unpaired and/or paired odors with an interval of 3-5 minutes between exposures. Odor exposure sequence was determined randomly. All recordings were performed at 18.5°C room temperature and 59% humidity.

Data were registered, processed, and extracted using a Matlab GUI developed by C. Deister, Brown University (https://github.com/cdeister/imageAnalysis_gui). Volumetric calcium imaging files (.raw) were adjusted to account for 26(z) slices and maximum Z-projections were made across time to collapse flies to a single Z-plane and converted to.tiff files. Images were then aligned and registered in X-Y using a total mean fluorescence as a template. ROIs were constructed from the neuropil using standard deviation and mean fluorescence templates to identify regions pertaining to VUMa3 neurons and then subsequently segmented to create the ROIs. Fluorescence values were extracted from identified ROIs and ΔF/F_o_ measurements were created using a moving-average of 5 frames to calculate the baseline fluorescence (F_o_). Fluorescence traces were Z-score normalized to their pre-stimulus air baseline (12 frames of exposure for the retrieval experiments). Average fluorescence traces across flies (n = 8-14) were visualized using ggplot2 in R Studio.

For laser power experiments the baseline fluorescence to air exposures were recorded at each of the different laser power intensities: 10, 12, 15, 20, and 25 mW. There was a 2–5 minute interval in between individual laser power baseline recordings and laser power order used was randomized within each fly. ROIs for the baseline fluorescence signal were segmented similarly as detailed above and were representative of large axons of the VUMa3 neuron that are present in the superior medial protocerebrum of the adult *Drosophila* brain. Additionally, for these baseline experiments a background ROI was placed in a region near the axons that is deemed to be representative of the rest of the background fluorescence. Representative ROIs from an example recording are schematized and detailed in Figure 3A.

For *in vivo* memory retrieval calcium imaging experiments MB021B-GAL4>UAS-GCaMP7b flies were conditioned to associate an odor cue with an intoxicating dose of ethanol and exposed the following day to both the paired stimulus (CS+) and unpaired stimulus (CS-) for 30 seconds while satiated or food-deprived (Figure 3D). When comparing the mean stimulus response, we defined a timepoint after stimulus onset and took the average normalized response across that time, comparing their responses within flies, across stimuli (Figure 3E, 3F).

### Connectomics

Reconstructions from FlyWire are based on EM image data that was released by Zheng et al 2018^169^ under a CC-BY-NC 4.0 International license. Data generated by FlyWire will be made publicly available to the scientific community, in accordance with FlyWire Principles. Synapse information is imported from the work of Buhman et al., 2021^170^, with synaptic “cleft_score” information incorporated into FlyWire from Heinrich et al 2018^171,172^. Neurotransmitter predictions are made from Eckstein et al 2024^169,172,173^, with infrastructure contributed by Davi Bock, Gregory Jefferis and Eric Perlman. Synapse information used in Figure 4C are retrieved and constructed using greater than 10 synapses for identification of synaptic targets with the braincircuits.io applications (Gerhard, Stephan, Lee, Wei-Chung Allen, Wilson, Rachel. (2022-2023). The BrainCircuits.io platform. https://braincircuits.io/. Manuscript in preparation) and identifying the top pre– and post-synaptic targets using the FlyWire connectivity application (https://prod.flywire-daf.com/dash/datastack/flywire_fafb_production/apps/fly_connectivity/) and FlyWire Codex database version 630 (https://codex.flywire.ai/).

### Data Visualization

All barplots, boxplots, linegraphs, piecharts throughout the manuscript were constructed using ggplot2 in RStudio (RStudio and R Binary Version 3.5)^174^. Heatmaps were constructed using standard built-in heatmap functions within MATLAB (Version R2019a (9.6.0.1072779)).

### Statistical Analysis

Parametric analyses were used unless assumptions of homogeneity of variance (Levene’s Test, P > 0.05), equal sample sizes, or normal distribution (Shapiro– Wilk normality test*, P* > 0.05) were violated. Grubb’s single outlier tests performed prior to statistical comparisons with p value of <0.01 were excluded from behavioral data. For parametric analyses the standard Student’s t-test, two-tailed, were performed to compare differences between two groups. One-way ANOVA (F > 0.05) was used for looking at the behavioral differences across 3 or more groups, followed by Tukey’s HSD and Holm’s inference statistic for planned contrasts between each genetic control (GAL4/+ and +/UAS-Shibire^ts1^) and the experimental group (GAL4>UAS-Shibire^ts1^). If assumptions were violated, non-parametric statistical tests consisted of Kruskal-Wallis/Wilcoxon tests with planned comparisons, same comparisons as those listed above for parametric analyses. Statistical analysis was performed using JMP software (JMP® Pro 15.2.0) where statistical significance is described as P < 0.05. All data is presented in graphs and figures as mean ± s.e.m. For behavior figures, individual data points are overlaid on the plots. One data point, or N equals about 60 flies, comprising the average preference indexes of two reciprocally trained odor groups. All analyses for calcium imaging experiments following a similar statistical pipeline. For retrieval experiments, since the same fly was exposed to the unpaired and paired odor at different times, a matched paired T-test was run using a specific stimulus response window indicated 5 seconds following stimulus onset. We used repeated ANOVA measures for the laser power baseline experiments since the same sample was measured at multiple laser power intensities. Follow-up statistical analysis tested planned contrasts between the two conditions (food-deprived and satiated) at each laser power intensity.

## SUPPLEMENTAL MATERIALS

**Supplementary Figure 1.**
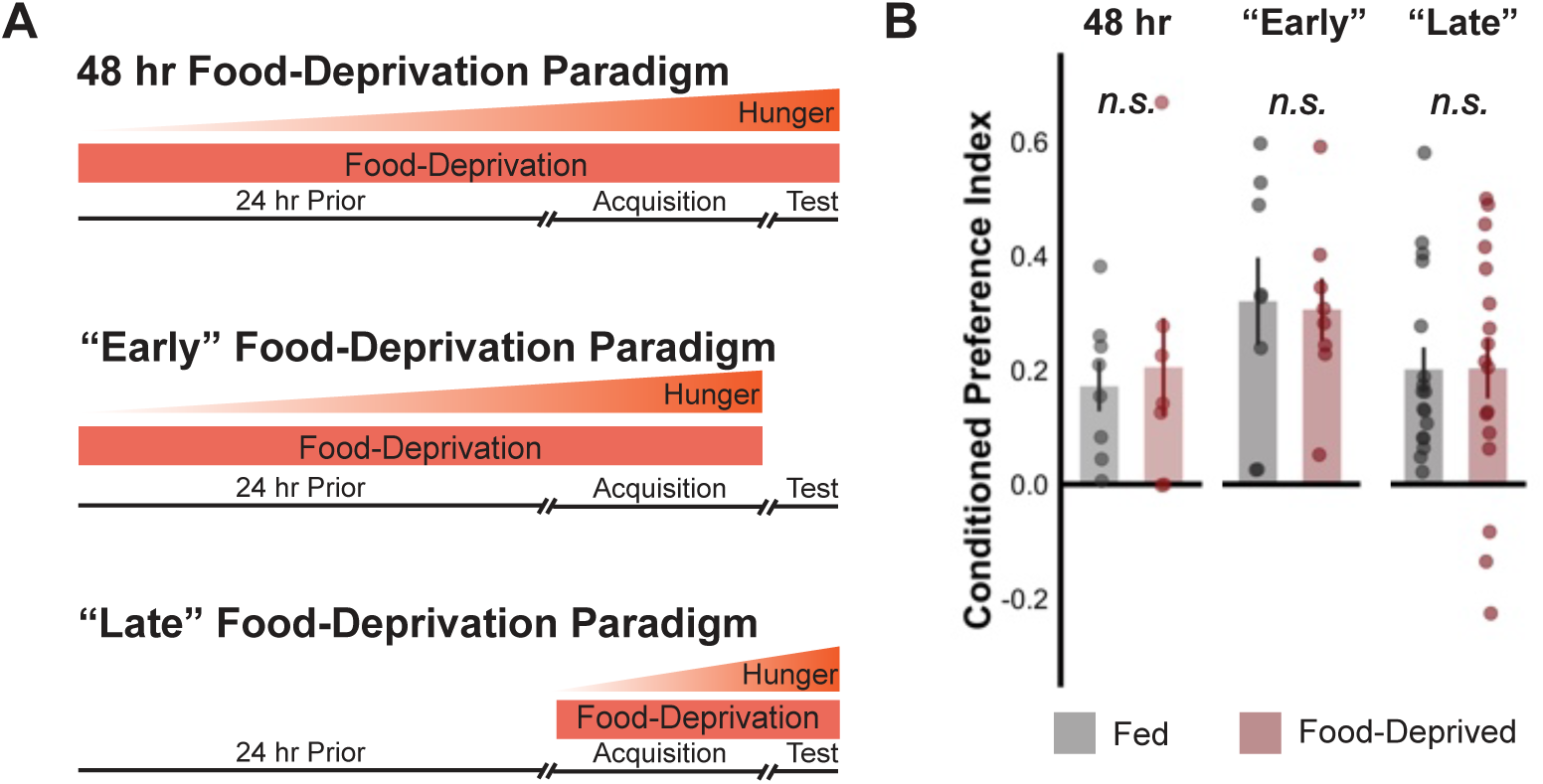
Food-deprivation does not impact ethanol preference in wild-type Canton S control flies. (A) Schematic of different food-deprivation paradigms used while performing ethanol conditioning. For the 48-hour food deprivation paradigm, the flies are placed on 2% agar vials 24 hours prior to the start of training and kept on agar throughout until the test period. The “early” food-deprivation paradigm similarly places the flies on agar 24 hours prior to the start of training, but are fed yeast pellets following the training/acquisition period. The “late” food-deprivation paradigm, food-deprives the flies from the start of training throughout training. (B) Regardless of the food-deprivation paradigm, food-deprivation does not alter conditioned ethanol preference when compared to satiated controls. Bar graphs illustrate mean +/-standard error of the mean. Raw data are overlaid on bar graphs. Each dot is an n of 1, which equals approximately 60 flies (30 per odor pairing).

**Supplementary Figure 2.**
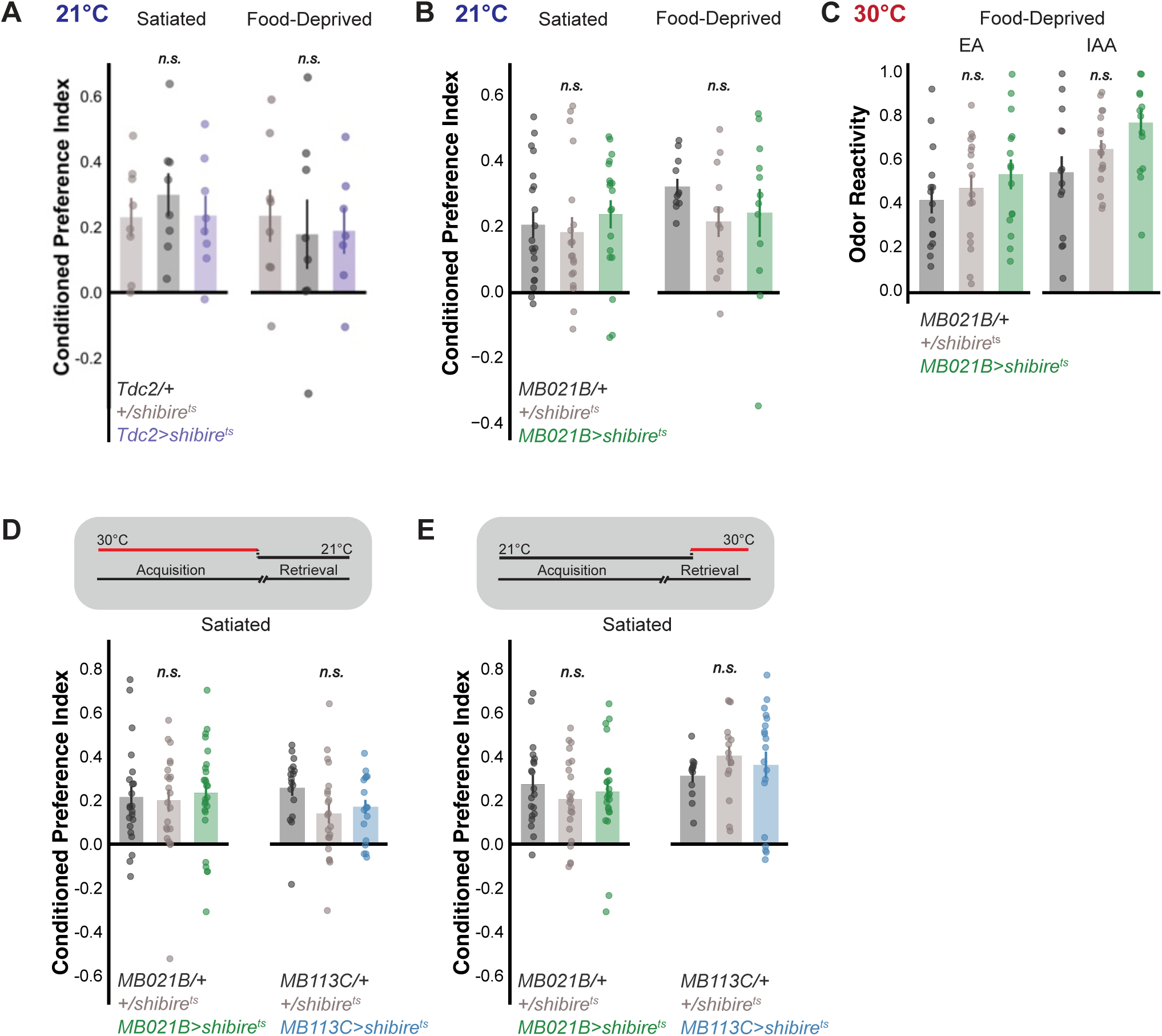
Tdc2-GAL4 and MB021B-splitGAL4 temperature controls. (A) Temperature controls for Tdc2-GA/.4 experiments, showing that there is no effect when these lines are at the permissive temperature for shibirets1, regardless of satiety state. (B) Temperature controls of MB021B-splitGA/.4 show no effect on ‘food-deprived’ ethanol preference. (xhibiting that the effect is a result of inactivating these neurons and not due to the introduction of transgenes. (C) Inactivation of MB021B-splitGA/.4 neurons and food depriving the flies shows no impact on odor reactivity to either odor, when compared to their genetic controls. (D) Thermogenetic inactivation of either MB021B or MB113C split-GA/.4 during the acquisition period does not result in a reduction of ethanol preference relative to transgenic controls when flies are satiated. (() Thermogenetic inactivation of either MB021B or MB113C split-GA/.4 during the retrieval period does not result in a reduction of ethanol preference relative to transgenic controls when flies are satiated. Bar graphs illustrate mean +/-standard error of the mean. Raw data are overlaid on bar graphs. (ach dot is an n of 1, which equals approximately 60 flies (30 per odor pairing or two odor reactivity scores).

**Supplementary Figure 3.**
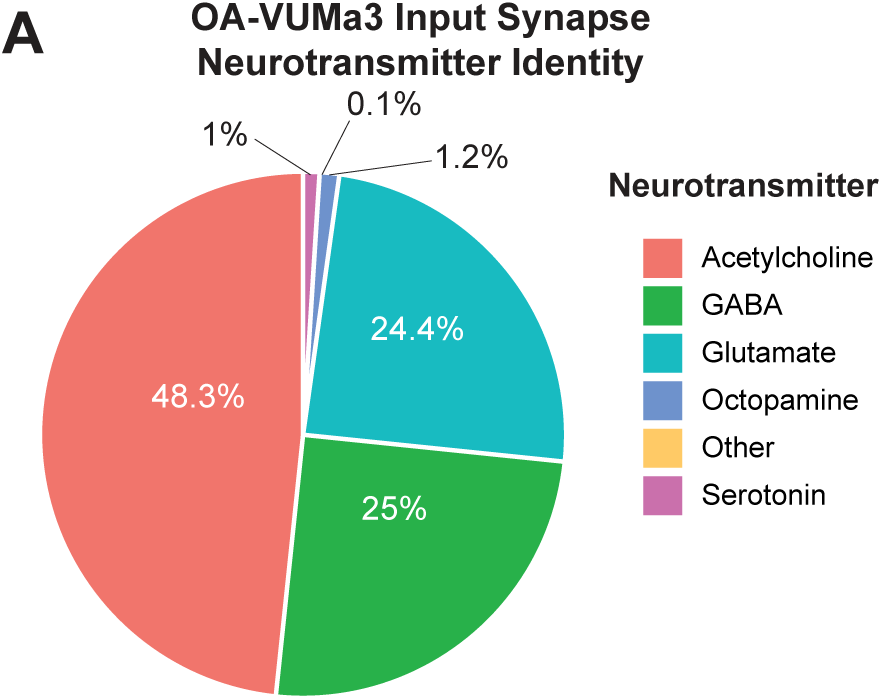
Predicted Neurotransmitter Identity of Pre-synaptic Inputs onto OA-VUMa3. (A) The pie chart depicts predicted input synapses onto OA-VUMa3 according to the FlyWire database. Acetylcholine is predicted to provide the most input onto VUMa3 with 48.3%, GABA at 25%, glutamate at 24.4%, octopamine at 1.2%, serotonin at 1%, and ‘Other’ at 0.1%.

**Supplemental Figure 4.**
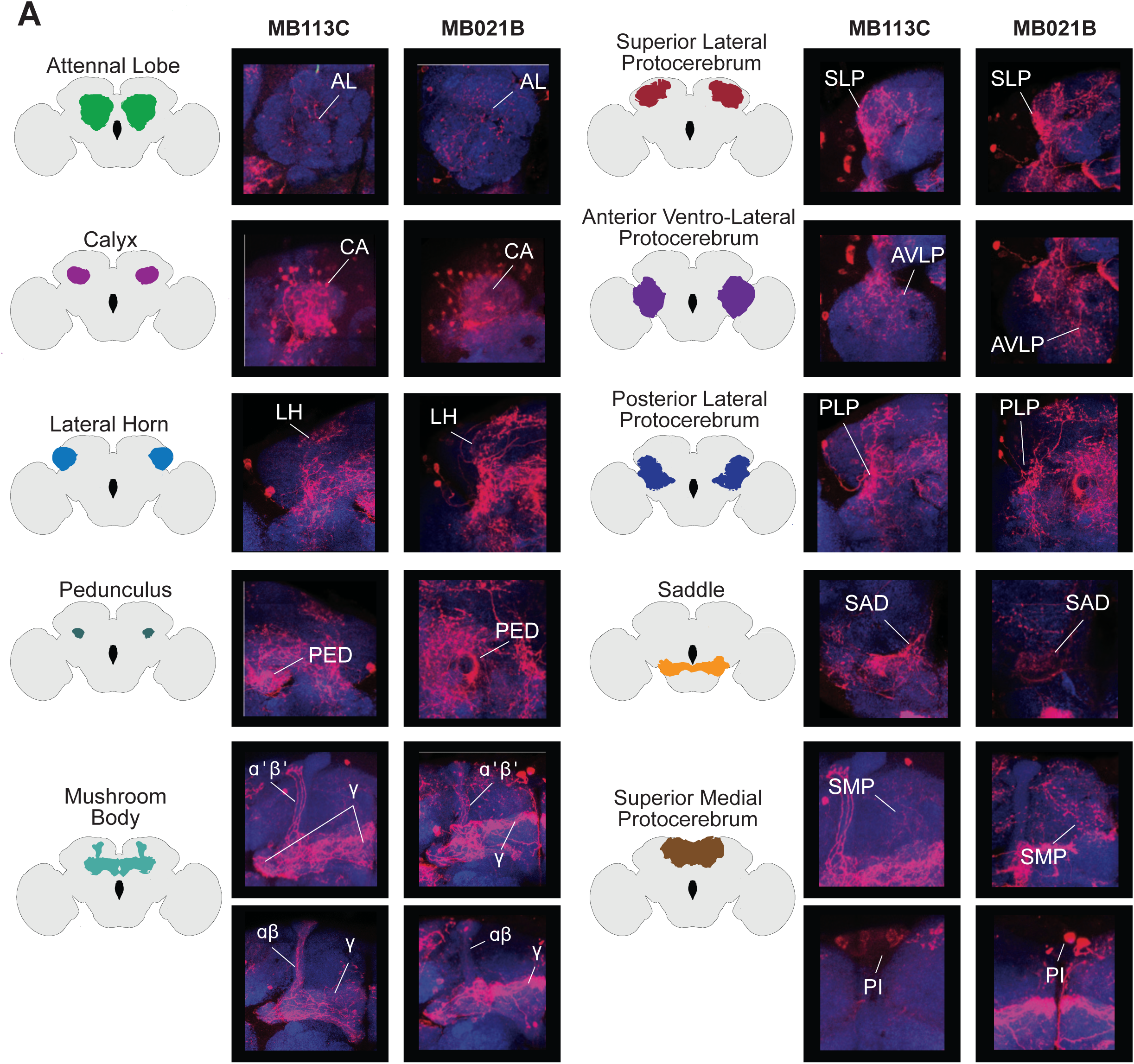
Post-synaptic targets of MB113C and MB021B. (A) Comparisons are displayed between the post-synaptic targets of MB113C-splitGAL4>trans-Tango and MB021B-splitGAL4>trans-Tango with region specific focuses throughout the brain, specified by the brain region schematics (left). Similarities in post-synaptic targets between MB113C and MB021B are displayed across regions of interest within the adult Drosophila brain. Both MB113C and MB021B innervate: neurons within the antennal lobe (AL), calyx (CA), lateral horn (LH), peduncle (PED), all of the Mushroom Body lobes, superior lateral protocerebrum (SLP), anterior ventrolateral protocerebrum (AVLP), posterior lateral protocerebrum (PLP), saddle (SAD), and superior medial protocerebrum (SMP). These similarities in innervation suggest that these post-synaptic targets may be of VPM4 and not VUMa3.

**Supplementary Figure 5.**
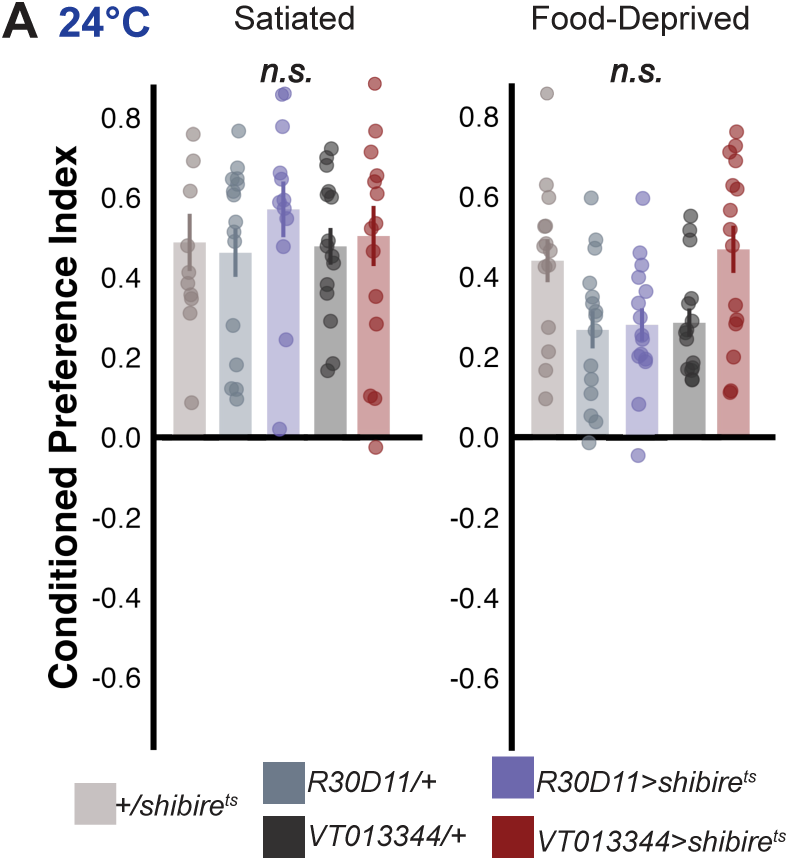
R30D11-GAL4 and VT013344-GAL4 temperature controls. (A) Temperature controls of R30D11-GAL4 and VT013344-GAL4 show no effect on ‘satiated’ or ‘food-deprived’ ethanol preference. Exhibiting that the effect is a result of inactivating these neurons and not due to the introduction of transgenes. Bar graphs illustrate mean +/-standard error of the mean. Raw data are overlaid on bar graphs. Each dot is an n of 1, which equals approximately 60 flies (30 per odor pairing).

